# Omicron-specific ultra-potent SARS-CoV-2 neutralizing antibodies targeting the N1/N2 loop of Spike N-terminal domain

**DOI:** 10.1101/2024.07.10.602843

**Authors:** Xiao Niu, Zhiqiang Li, Jing Wang, Fanchong Jian, Yuanling Yu, Weiliang Song, Ayijiang Yisimayi, Shuo Du, Zhiying Zhang, Qianran Wang, Jing Wang, Ran An, Yao Wang, Peng Wang, Haiyan Sun, Lingling Yu, Sijie Yang, Tianhe Xiao, Qingqing Gu, Fei Shao, Youchun Wang, Junyu Xiao, Yunlong Cao

**Author notes:** Corresponding author (JX); (YC). These authors contributed equally (Xiao Niu, Zhiqiang Li, Jing Wang).

## Abstract

A multitude of functional mutations continue to emerge on the N-terminal domain (NTD) of the spike protein in SARS-CoV-2 Omicron subvariants. Understanding the immunogenicity of Omicron NTD and the properties of antibodies elicited by it is crucial for comprehending the impact of NTD mutations on viral fitness and guiding vaccine design. In this study, we find that most of NTD-targeting antibodies isolated from individuals with BA.5/BF.7 breakthrough infection (BTI) are ancestral (wildtype or WT)-reactive and non-neutralizing. Surprisingly, we identified five ultra-potent neutralizing antibodies (NAbs) that can only bind to Omicron but not WT NTD. Structural analysis revealed that they bind to a unique epitope on the N1/N2 loop of NTD and interact with the receptor-binding domain (RBD) via the light chain. These Omicron-specific NAbs achieve neutralization through ACE2 competition and blockage of ACE2-mediated S1 shedding. However, BA.2.86 and BA.2.87.1, which carry insertions or deletions on the N1/N2 loop, can evade these antibodies. Together, we provided a detailed map of the NTD-targeting antibody repertoire in the post-Omicron era, demonstrating their vulnerability to NTD mutations enabled by its evolutionary flexibility, despite their potent neutralization. These results highlighted the importance of considering the immunogenicity of NTD in vaccine design.

**Author Summary:** COVID-19 pandemic caused by severe acute respiratory syndrome coronavirus 2 (SARS-CoV-2) continues to be a major global public health concern four years after its emergence. The N-terminal domain (NTD) is a critical component of the spike glycoprotein, which is pivotal for SARS-CoV-2 cellular entry and serves as a primary target for antibody therapeutics and vaccine development. Characterizing the properties of antibodies elicited by NTD of Omicron sublineages is crucial for understanding viral evolution and guiding vaccine design. Here, we show that Omicron infection after vaccination induces majorly non-neutralizing NTD antibodies. Still, we identified a class of ultra-potent neutralizing antibodies (NAbs) which specifically bind to the NTD of Omicron sublineages. These NAbs neutralize the virus by competing with ACE2 and blocking ACE2-mediated S1 shedding. Structural analyses reveal that these antibodies target a unique epitope on the N1/N2 loop of NTD, and intriguingly interact with the receptor-binding domain (RBD) of spike glycoprotein. This class of NAbs with the special binding pattern, are escaped by BA.2.86 and BA.2.87.1 sublineages, shedding light on the role of recently emerged mutations in the N1/N2 loop of NTD. Our findings provide fresh insights into the immunogenicity of Omicron NTD, highlighting its capacity for antibody evasion due to its evolutionary flexibility. This underscores the importance of carefully considering the NTD component in vaccine design.

## Introduction

Recently emerging Omicron subvariants, such as HK.3, BA.2.86, JN.1, BA.2.87.1, KP.2 and KP.3, have shown significant evasion capabilities against the humoral immunity elicited by Omicron infection or vaccination [1–3]. Convergent mutations on the spike protein of these lineages are considered to be responsible for their escape. While epitope mapping of antibodies targeting the RBD has enabled prediction of RBD evolution [4–8], the evolutionary pressure caused by NAbs targeting non-RBD regions, especially the NTD of the spike protein, are largely unclear.

NTD has been demonstrated to be immunogenic in multiple pre-Omicron studies [9, 10]. NAbs induced by NTD also have been shown responsible for the cross-neutralization against BA.5 in BA.2 BTI convalescents [11]. Concurrently, NTD continues to evolve, notably through insertions and deletions of multiple amino acids, diverging from RBD mutations [12–15]. Within Omicron subvariants, BA.2 exhibited distinct NTD mutations compared to BA.1, which has the notable HV69-70del and 143-145del. Subsequently, K147E and W152R emerged in BA.2.75, 69-70del re-emerged in BA.5, Y144del re-appeared in the XBB sublineage, Q52H emerged in EG.5.1 [16], ins16MPLF appeared in BA.2.86/JN.1 [17, 18], and long deletions at positions 15-26 and 136-146 appeared in BA.2.87.1 [19]. Most recently, Q183H and S31del were identified in the prevailing strain LB.1, KP.2.3 [20] and XDY. Some of these mutations have been linked to enhanced viral fitness in various variants [21], including HV69-70del for viral infectivity [22], Y144del or K147E+W152R for NTD NAb evasion [4, 23], and S31del for increased infectivity and immune evasion [20]. However, the functional implications of these extensive mutations, particularly those occurring subsequent to XBB, as well as their potential to facilitate escape from NAbs targeting corresponding epitopes, remain to be elucidated.

Given that current mRNA vaccines against SARS-CoV-2 are primarily based on the full-length spike protein, the immunogenicity of NTD must be considered in vaccine design strategies. A comprehensive investigation into the immunogenicity of Omicron NTD and characterization of Omicron induced NTD antibodies would facilitate the prediction of NTD mutations. Some NTD epitopes and their elicited antibodies have been characterized [9, 24–29]. The majority of existing NTD-targeting NAbs with notable potency recognize an antigenic supersite (residues 14–20, 140– 158, and 245–264) [10, 27, 28, 30], which was disrupted by mutations on NTD in XBB sublineages. Therefore, it is imperative to examine the NTD-targeting antibody repertoire induced by Omicron BTIs to identify potential NAbs and determine their binding epitopes.

Here, we isolated 361 NTD-binding monoclonal antibodies from individuals with BA.5/BF.7 BTI, and five of them exhibited exceptionally potent neutralizing capabilities against Omicron subvariants, including XBB.1.5 and HK.3.1. We found that this class of antibodies can compete with ACE2 albeit to a lesser degree and inhibit ACE2-mediated S1 shedding, which may be the mechanisms of neutralization. Structural analysis using cryogenic electron microscopy (cryo-EM) reveals that these antibodies target a novel NTD epitope on the N1 and N2 loop, distinct from the supersite previously identified [10, 27, 28, 30]. Interestingly, they can establish hydrogen bonds with RBD through their light chain, thereby maintaining a certain level of RBD affinity. Deep mutational scanning (DMS) data indicate that currently circulating variants have not developed single-point substitutions capable of evading these antibodies. However, BA.2.86 and BA.2.87.1 escaped these antibodies through 16MPLF insertion or 15-23 deletion on their NTDs, respectively, suggesting that the flexibility of NTD enables the virus to evolve atypical and radical mutations to circumvent neutralization by NTD-targeting antibodies.

## Results

### Omicron infection induces non-neutralizing or easily evadable antibodies

To investigate the NTD antibody repertoire induced by Omicron infection, we collected blood samples from 31 individuals who had received 2-3 doses of CoronaVac before being infected by BA.5, and 48 individuals who had received 3 doses of CoronaVac prior to BF.7 infection (S1A Fig; S1 Table).

Then, NTD^WT/BA.5^-positive memory B cells (CD20^+^, CD27^+^, IgM/IgD^-^) were isolated from peripheral blood mononuclear cells (PBMCs) of these convalescents using fluorescence-activated cell sorting (FACS) (S1B Fig). Antibody sequences are determined by single-cell V(D)J sequencing, as described previously [5–7]. A total of 361 mAbs were finally expressed and purified as human IgG1 (S2 Table). Binding capabilities of the isolated mAbs against NTD^WT^, NTD^BA.2.75^, NTD^BA.5^ and NTD^XBB^ were determined by enzyme-linked immunosorbent assays (ELISA), and their neutralizing activities against D614G, BA.2, BA.5, BA.2.75, BQ.1.1, BQ.1.1+Y144del, XBB, XBB.1 and XBB.1.5 were measured via pseudovirus neutralization assays (Figs 1A and S1C).

**Fig 1.**
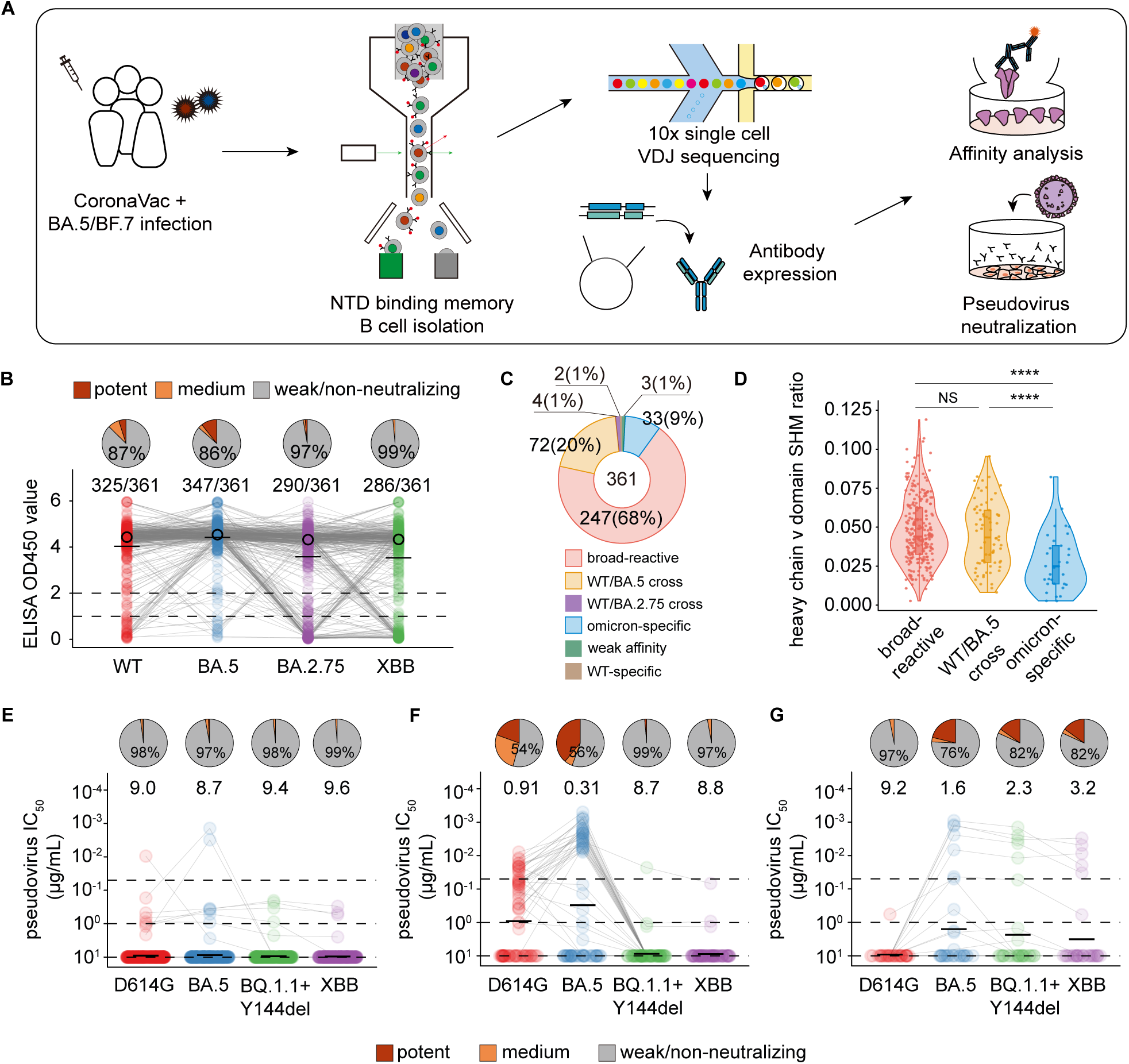
Omicron infection induces Omicron-specific NTD targeting monoclonal antibodies. (A) Workflow of this study. (B) Binding affinity (ELISA OD450) and pseudovirus-neutralization activities(Half maximal inhibitory concentration, IC50) of isolated monoclonal antibodies (n = 361) against WT, BA.5, BA.2.75 and XBB. Medians and means are depicted as black circles or short lines in the figure. Pie charts represent the distribution of pseudovirus-neutralization activities of high-affinity antibodies (OD450 > 2) against the corresponding variant. Neutralization potency is categorized as potent (IC_50_ ≤ 0.05), medium (0.05 < IC_50_ ≤ 1), and weak/non-neutralizing (IC_50_ > 1). (C) Categories of the 361 antibodies based on affinity. Define high affinity for OD450 ≥ 2 and low affinity for OD450 < 2. The antibodies are categorized into six groups: Broad-reactive, showing high affinity to WT, BA.2.75, BA.5, and XBB NTDs (247 of 361); WT/BA.5 cross-reactive, showing high affinity to WT and BA.5 NTDs (72 of 361); WT/BA.2.75 cross-reactive, showing high affinity to WT and BA.2.75 NTDs (4 of 361); Omicron-specific, showing high affinity to BA.2.75, BA.5, or XBB NTDs but low affinity to WT NTD (33 of 361); Weak affinity, showing low affinity to WT, BA.2.75, BA.5, and XBB NTDs (3 of 361); and WT-specific, showing high affinity to WT NTD but low affinity to BA.2.75, BA.5, and XBB NTDs (2 of 361). (D) The heavy chain V domain somatic hypermutation (SHM) ratio of broad-reactive, WT/BA.5 cross and Omicron-specific antibodies. Two-tailed Wilcoxon rank-sum tests was used to examine the significant differences of unpaired samples. *P < 0.05, **P < 0.01, ***P < 0.001, ****P < 0.0001; NS, not significant (P > 0.05). (E-G) Pseudovirus-neutralization activities of broad-reactive (E) WT/BA.5 cross (F) and Omicron-specific (G) antibodies. Geometric mean titres (GMTs) are shown.

All 361 antibodies exhibited binding affinity to either WT, BA.5, BA.2.75 or XBB NTD (OD450> 1) (Figs 1B and S1C). Interestingly, antibodies with high affinity (OD450 > 2) are predominantly weakly neutralizing or non-neutralizing (IC_50_ > 1) against the corresponding variant (Fig 1B), suggesting a large proportion of NTD-targeting but non-neutralizing antibodies induced by Omicron infection.

Based on the affinity to NTDs (ELISA OD450 values), the 361 antibodies were categorized into 6 groups (Fig 1C). The majority of the antibodies (68%) are broad-reactive, binding to conserved epitopes that remain unmutated across the four variants. WT/BA.5 cross-reactive antibodies (20%) can bind to both WT and BA.5 but are escaped by BA.2.75 and XBB, while Omicron-specific antibodies (9%) are unable to bind to WT and only bind to Omicron NTD. Broad-reactive and WT/BA.5 cross-reactive antibodies exhibit significantly higher levels of somatic hypermutation (SHM) than Omicron-specific antibodies (Fig 1D), indicating that these two categories of antibodies were elicited during vaccination, recalled and further matured during Omicron BTI, whereas Omicron-specific antibodies are newly generated in response to BA.5/BF.7 BTI.

Subsequently, we evaluated the pseudovirus neutralization profiles of these three categories of antibodies. Over 97% of broad-reactive antibodies demonstrated weak/non-neutralizing activity (IC_50_>1) against D614G, BA.5, BQ.1.1+Y144del and XBB (Fig 1E), despite their high affinity for all four NTDs. This suggests that they target conserved, but non-neutralizing epitopes. Given their weak/non-neutralizing nature, these antibodies do not exert immune pressure on the virus, thus are not escaped and repeatedly recalled during infections.

Existing studies have indicated that neutralization efficacy of mAbs targeting the NTD supersite like 4A8 can be eliminated by Y144del [31]. Additionally, we have previously reported the significant reduction in neutralizing titers of BA.5 BTI convalescent plasma against emerging SARS-CoV-2 variants carrying convergent mutations on RBD, and we found that BQ.1 and BQ.1.1 only moderately evade BA.5 BTI plasma, while BQ.1.1+Y144del, XBB and BA.2.75 sublineages exhibited dramatic evasion against them, indicating a non-negligible contribution from supersite-targeting NAbs [6]. WT/BA.5 cross-reactive antibodies exhibited higher neutralization titers against D614G and BA.5, with 40∼50% showing potent or moderate neutralization capacity. However, nearly all of them were escaped by BQ.1.1+Y144del or XBB (Fig 1F). Moreover, most of antibodies with a BA.5 IC_50_ < 10 utilized the IGHV1-24 gene (S2 Table), which was frequently utilized by supersite-targeting antibodies [32]. This indicates that BA.5/BF.7 BTI elicits a significant number of supersite-targeting antibodies capable of neutralizing BA.5. However, they are readily escaped by subsequent variants.

Remarkably, among the Omicron-specific antibodies, about 16% (5 of 33) demonstrated robust and ultra-potent neutralizing activity across all tested Omicron variants (Fig 1G). These antibodies likely emerged post-Omicron infection (Fig 1D), potentially targeting novel epitopes specific to Omicron that have remained intact in XBB.

### A class of ultra-potent Omicron-specific anti-NTD NAbs

To further characterize these five antibodies, their binding affinity was first verified using surface plasmon resonance (SPR). The results revealed that they exhibited high affinity for BA.5 NTD, XBB NTD, and XBB.1.5 spike, particularly for the XBB.1.5 spike, with KD values reaching the order of 10^-10 (S2 Fig). Pseudovirus neutralization assays were then performed using recently prevalent variants with high evasion capability, including XBB.1.5, EG.5.1, and HK.3.1 (Fig 2A). The antibodies demonstrated exceptional neutralizing abilities against all tested Omicron pseudoviruses, with IC_50_ values below 0.01 or even lower. Subsequently, we analyzed and compared the VDJ sequences and binding epitopes of these antibodies. Three of them utilize the identical heavy chain variable gene segment, IGHV1-18, and two share a common light chain variable gene segment, IGLV6-57 (Fig 2A). Multiple sequence alignment (MSA) revealed a notable sequence similarity within the heavy chain complementarity-determining region 3 (CDR3) among these antibodies, compared to other antibodies within the same batch (S3A Fig). Additionally, the five NAbs demonstrated high competition levels in competitive SPR experiments (Fig 2B), indicating that they target overlapping NTD epitopes. Their epitopes also appear to be distinct from the WT/BA.5 cross-reactive antibodies BD57-0617 and BD57-0740 (they are also supersite-targeting antibodies, see S3B Fig), the broad-reactive antibodies BD57-0536 and BD57-0569, and the previously reported NTD antibodies C1717 (group III) and C1520 (group IV) [10].

**Fig 2.**
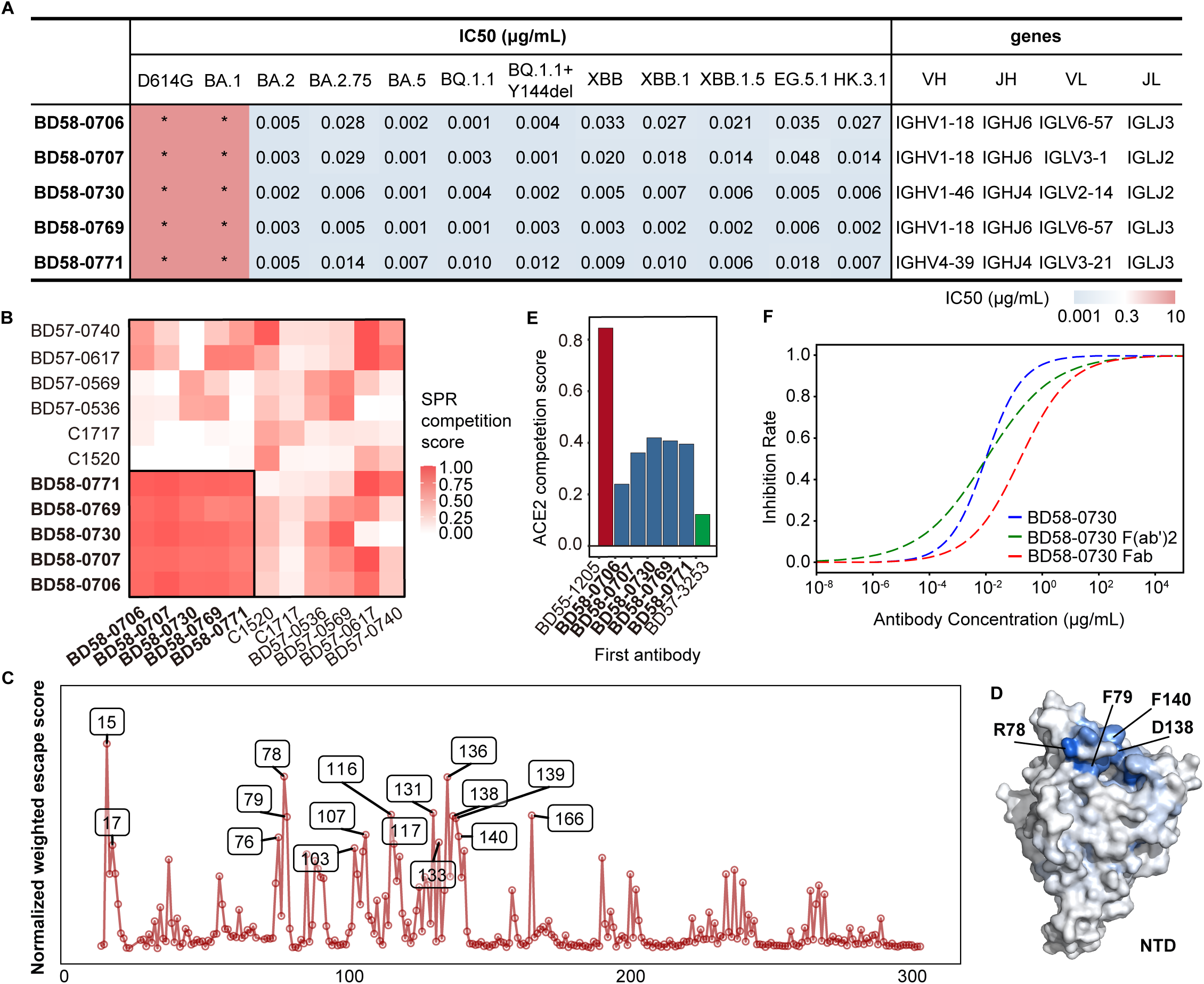
A class of ultra-potent Omicron-specific NTD-targeting monoclonal antibodies. (A) Pseudovirus-neutralizing IC_50_ of 5 ultra-potent NTD-targeting NAbs against SARS-CoV-2 D614G, BA.1, BA.2, BA.2.75, BA.5, BQ.1.1, BQ.1.1+Y144del, XBB, XBB.1, XBB.1.5, EG.5.1 and HK.3.1, and their usage of heavy chain V (VH), heavy chain J (VJ), light chain V (VL) and light chain J (VJ) genes. (B) Heatmap for pair-wised SPR competition scores of 5 ultra-potent NTD-targeting NAbs and BD57-0617, BD57-0740, BD57-0536, BD57-0569, C1717 and C1520 on XBB.1.5 spike. (C) Calculation of immune pressure on each NTD site (residues 13-304 in WT index) and mutation of BD58-0730 escape mutation profiles weighted by BA.5. (D) Average DMS site escape scores of BD58-0730 are indicated on the structure model of the SARS-CoV-2 BA.5 NTD (PDB: 7XNQ). Hotspot residues are indicated by arrows. (E) SPR competition scores of 5 ultra-potent NTD-targeting NAbs, BD57-3253 and BD55-1205 as first antibody against ACE2. (F) Pseudovirus-neutralization curve of full-length IgG, Fab and F(ab’)2 forms of BD58-0730. Data for each form were obtained from a representative neutralization experiment.

To investigate mutations that could potentially escape these antibodies, and further characterize the epitopes targeted by these antibodies, we also constructed duplicated deep mutational scanning (DMS) libraries based on BA.5 NTD, as we previously reported [6]. Through DMS, we profiled the mutation escape map of BD58-0730 (Fig 2C) and projected the results onto the NTD structural model (Fig 2D). Consistent with competition SPR experiments, the escape epitopes for BD58-0730 do not include Y144; instead, they are concentrated within the N1 (15-20) and N2 (69-77) loops. Moreover, they exhibited a certain degree of competition with ACE2 (Fig 2E). In comparison, the typical non-ACE2 competitive antibody BD57-3253[33] shows negligible competition, while the ACE2 mimicry antibody BD55-1205 [34] exhibit a competition score of 0.85 (Fig 2E).

Intriguingly, we observed that the Fab and F(ab’)2 forms of BD58-0730 exhibit a modestly reduced, yet still substantial neutralizing capacity against XBB.1.5 pseudovirus compared to the full-length IgG form (Fig 2F). The IC_50_ of its F(ab’)2 and Fab forms against XBB.1.5 are 0.0081 μg/ml and 0.12 μg/ml, respectively, compared to 0.0078 μg/ml for the full-length IgG form. Moreover, the F(ab’)2 demonstrated an ACE2 competition level on par with that of the full-length IgG (S3C Fig). Certain NTD NAbs have been demonstrated to require the intact IgG form to exert their neutralizing function, or they necessitate the assistance of Fc effector functions for optimal protection efficacy [25, 35]. Unlike these antibodies, the five antibodies identified in this study do not rely on the Fc region to exert their neutralizing capability, nor do the competition of these antibodies with ACE2 is achieved through steric hindrance by Fc. Under natural conditions, the binding of ACE2 to spike induces the shedding of the S1 glycoprotein [36]. ACE2-competitive antibodies targeting the NTD have been shown to be able to inhibit ACE2-induced S1 shedding [30], which was observed with these five antibodies (S4 Fig).

In conclusion, these antibodies represent a class of potent NAbs targeting similar epitopes specific to Omicron, which appear to achieve their neutralizing effects through mechanisms including ACE2 competition and blockage of ACE2-mediated S1 shedding.

### Cryo-EM structural analysis reveals that BD58-0730 targets N1/N2 loop of NTD

Nevertheless, these observations seem inadequate to fully account for the remarkably potent neutralizing capabilities exhibited by this class of antibodies. To further probe the molecular mechanisms underlying their binding affinity and neutralizing potency, we employed cryo-EM to resolve the structure of BD58-0730 in complex with the XBB.1.5 spike trimer (S5 Fig). The structural analysis revealed that BD58-0730 binds to the 1-up-2-down spike at a stoichiometry of 3:1 (Fig 3A).

**Fig 3.**
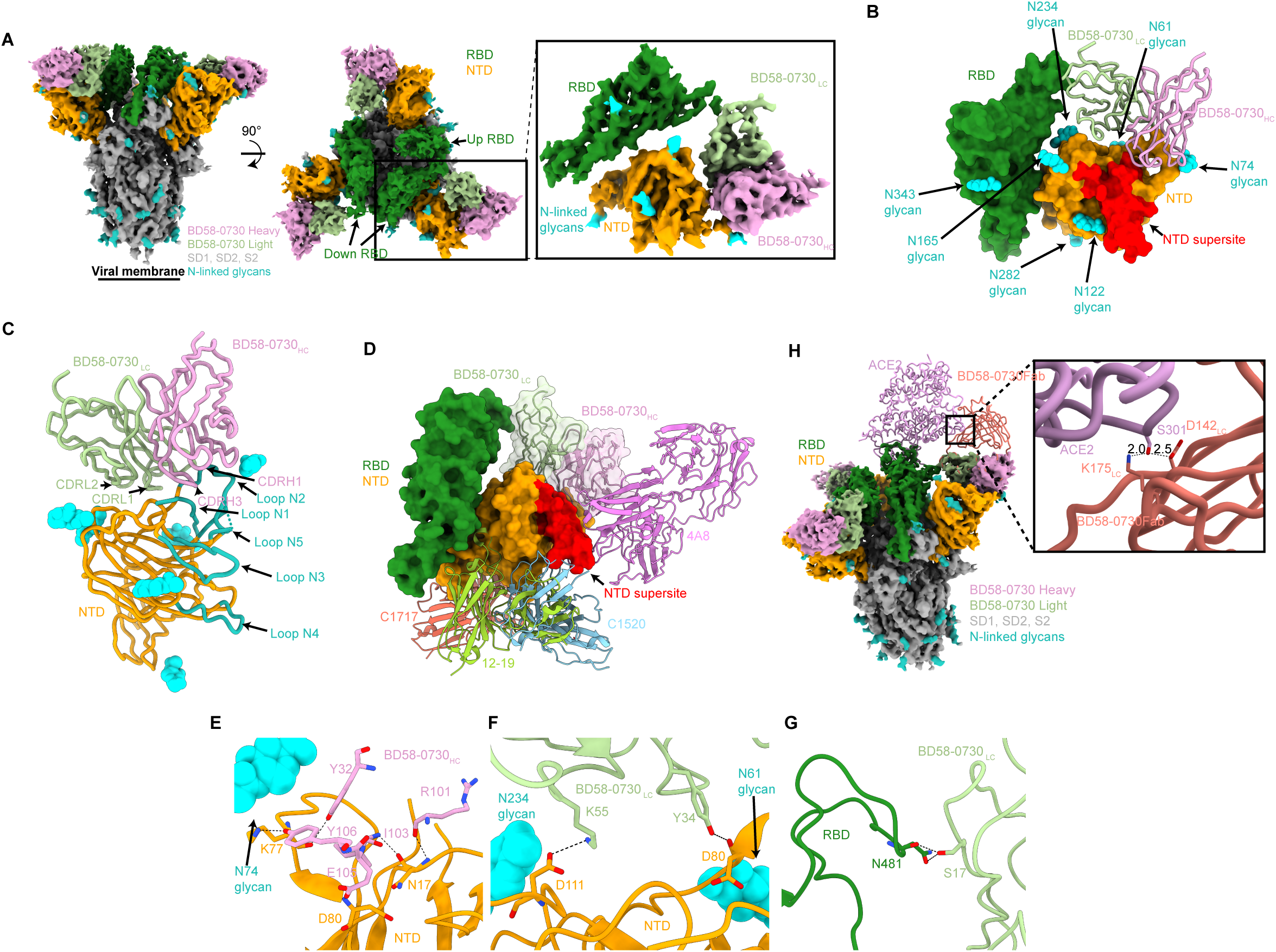
Cryo-EM structure suggests BD58-0730 targets a novel NTD epitope. (A) 3.55 Å cryo-EM density for BD58-0730-S 6P trimer complex. Inset: 3.77 Å locally refined cryo-EM density for BD58-0730 variable domains (shades of light green for light chain and pink for heavy chain) bound to NTD (orange) and RBD (green). (B) Close-up view of BD58-0730 variable domains (light green and pink licorice) binding to NTD (surface rendering, orange) and RBD (surface rendering, green). Supersite is highlighted on NTD for reference (red). (C) Close-up view 0f antibody BD58-0730 recognition of NTD, CDR H1, H3, L1, L2 marked out. NTD N1-5 loops colored in cyan. (D) Overlay of VH (pink) and VL (light green) domains of BD58-0730, C1717 (PDB: 7UAR; coral), 12-19 (PDB: 7UKM; yellow green), and C1520 (PDB: 7UAQ; cyan) and Fab of 4A8 (PDB:7C2L; magenta) after alignment on NTD residues 27–303, illustrating distinct binding poses to NTD epitopes adjacent to the supersite (red). (E-F) Residue-level contacts between the NTD (orange) with (E) BD58-0730 heavy chain (pink) (F) light chain (light green). Glycans are colored cyan. (G) Residue-level contacts between the RBD (green) with BD58-0730 light chain (light green). (H) Overlay of BD58-0730-S6P trimer complex, RBD-ACE2 complex (PDB:7C8D, purple) after alignment on up RBD, and BD58-0730 Fab (predicted by swiss model, tomato) alignment on VH and VL. Inset: closest distance between ACE2 and light chains.

The main recognition region of BD58-0730 is not located at the supersite but in N1 and N2 loops, especially N1 loop, surrounded by N61, N74, N165, and N234 glycans (Figs 3B and 3C), consistent with DMS results (Fig 2D). This may be attributable to the removal of N17 glycosylation by T19I mutation, which was frequently observed in Omicron subvariants, excluding BA.1. This removal creates a novel epitope within the NTD, enabling the binding of antibodies like BD58-0730 to the NTD. This also explains why such antibodies cannot neutralize BA.1 (Fig 2A).

Structural alignment reveals that the epitopes targeted by BD58-0730 do not overlap with those targeted by C1520, C1717 [10], 12-19 [30], and 4A8 [9] (Fig 3D). The heavy chain of BD58-0730 assumes a predominant role in the recognition of the antibody, with the buried surface area (BSA) of the NTD epitope on the heavy chain (HC) constituting ∼562 Å^2^ out of the total BSA of ∼933 Å^2^. Y32, R101, I103, E105, and Y106 on CDRH3 (97-111) and CDRH1 (26-33) have hydrogen bonding interactions with N17, K77, and D80 of NTD (Fig 3E). The light chain covers the remaining BSA of ∼371 Å^2^, where Y34 on CDHL1 and d80 on NTD have hydrogen bonding interactions, while K55 on CDRL2 and D111 on NTD have salt bridge interactions. (Fig 3F). Sequence analysis reveals an enrichment of tyrosine (Y) residues within the heavy chain CDR3 region of these five antibodies (S2 Table), suggesting a similar binding motif as Y106 in BD58-0730. The presence of Y34 in the light chain of BD58-0730 contributes to hydrogen bond formation with D80 of the NTD (Fig. 3F), and was found in other three of the antibodies excluding BD58-0771 (S2 Table).

Surprisingly, we observed a close hydrogen bond between S17 of BD58-0730’s light chain and N481 of the RBD (Fig 3G). Interestingly, the N481K mutation does not abrogate the neutralizing activity of these antibodies (S6 Fig), suggesting that their interaction with the RBD is not the sole mechanism underlying their neutralization ability. In the other four antibodies, position 17 is threonine (T). T and S differ by only one methylene group, possessing similar chemical properties, both capable of forming hydrogen bonds with hydroxyl groups on their side chains. Hence, T17 may serve a function analogous to S17. Structural prediction and alignment were employed to construct a conformational model in which both ACE2 and BD58-0730 are bound to the spike trimer (Fig 3H). The proximity between BD58-0730’s light chain and ACE2, at a distance of less than 3Å, implies potential steric clash impeding the interaction of the ’up’ conformation RBD with ACE2. This finding is consistent with SPR results where BD58-0730 demonstrated competitive binding with ACE2. Consistent with the epitopes targeted by BD58-0730, we hypothesize that mutations on the N1 or N2 loop of the NTD may confer evasion from these antibodies.

### BA.2.86 and BA.2.87.1 lineages escape NTD-targeting NAbs

The latest prevalent variants have undergone consecutive amino acid insertions or deletions in N1/N2 loop of NTD, which are likely to confer escape from antibody neutralization (Fig 4A). As expected, we found that all five antibodies were unable to neutralize BA.2.86 or BA.2.87.1 pseudovirus (Fig 4B).

**Fig 4.**
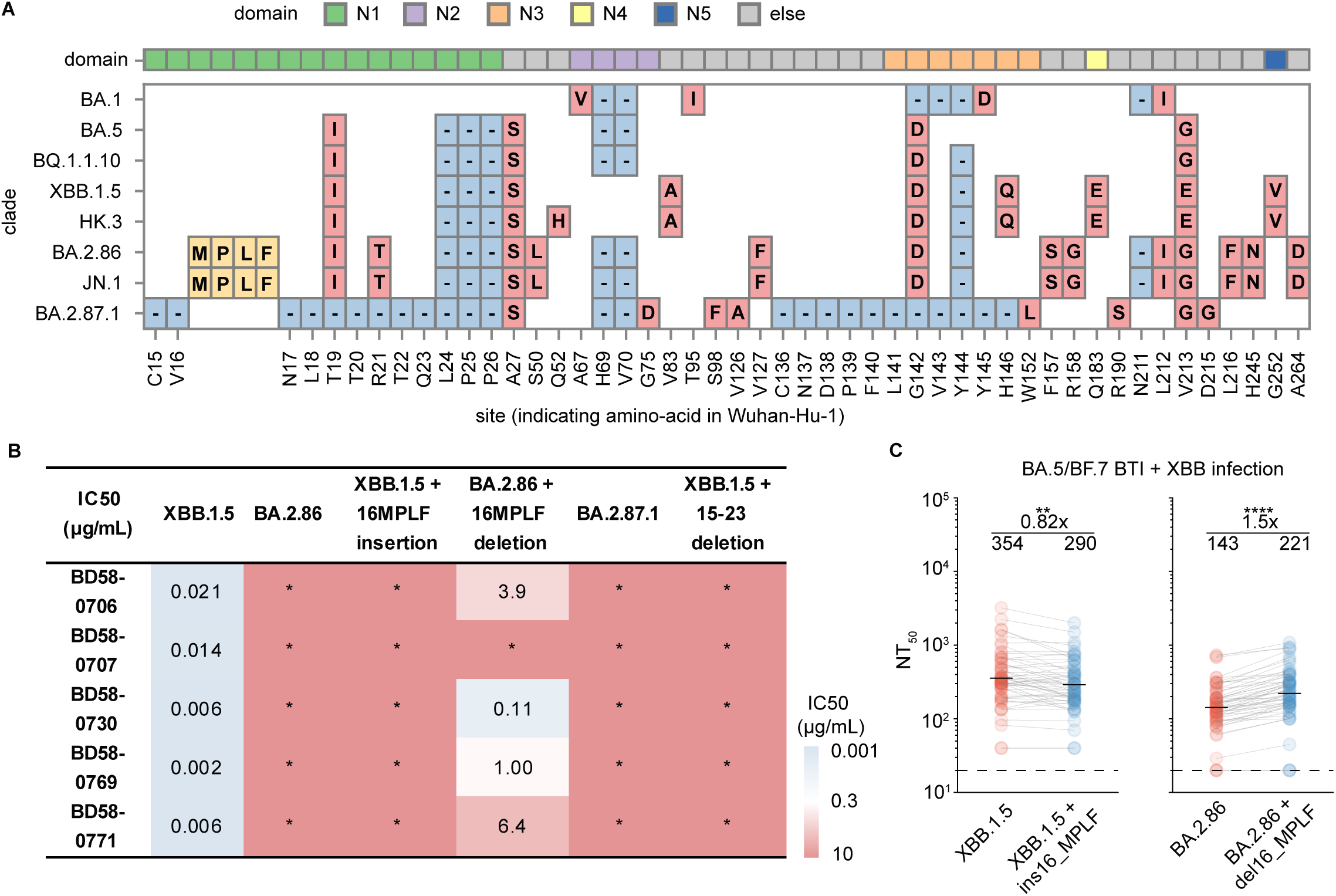
16MPLF insertion on BA.2.86 and 15-23 deletion on BA.2.87.1 NTD escape the five ultra-potent Omicron-specific antibodies. (A) NTD mutations carried by SARS-CoV-2 variants. N1-N5 loops are shown as green, purple, orange, yellow and blue. (B) Pseudovirus-neutralizing IC_50_ of 5 ultra-potent NTD-targeting NAbs against XBB.1.5, BA.2.86, XBB.1.5 + MPLFins, BA.2.86 + MPLFdel, BA.2.87.1 and XBB.1.5 + 15-23del. (C) 50% neutralization titers (NT_50_) against XBB.1.5, XBB.1.5 + MPLFins, BA.2.86 and BA.2.86 + MPLFdel of plasma samples from convalescents infected by XBB after 3 doses of CoronaVac (n = 54).

To further determine the mutations contributing to the escape of BA.2.86 and BA.2.87.1 from these antibodies, we constructed XBB.1.5 pseudoviruses carrying an insertion of 16MPLF (XBB.1.5+16MPLFins) or a deletion spanning amino acids 15-23 (XBB.1.5+15-23del). These antibodies all failed to neutralize XBB.1.5+16MPLFins or XBB.1.5+15-23del, but four of them could neutralize BA.2.86+16MPLFdel (Fig 4B). These results suggest that BA.2.86 and BA.2.87.1 evade neutralization by these antibodies through extensive insertion-deletion mutations.

To examine the role of such antibodies within the context of existing herd immunity, we analyzed convalescent plasma obtained from individuals who had experienced BTIs with BA.5/BF.7 and XBB (S1 Table). The 16_MPLF insertion on the background of XBB.1.5 caused a significant reduction in neutralizing potency against XBB.1.5, while the deletion of 16_MPLF based on BA.2.86 significantly enhanced the plasma neutralization titer compared to that against BA.2.86 (Fig 4C), indicating a substantial presence of such antibodies against current immune background. In summary, this class of Omicron-specific antibodies we isolated from individuals with Omicron BTI, targeting the N1/N2 loop of the NTD, significantly contributes to plasma neutralization. However, they are susceptible to escape due to the highly diverse mutations occurring within the NTD.

## Discussion

Newly emerged Omicron variants tend to show resistance to humoral immunity induced by previous infections and vaccinations, evading RBD-targeting NAbs through rapid mutation [1, 37]. Besides studies on RBD, there has also been research concerning the NTD since the onset of the COVID-19 pandemic. One study demonstrated that a class of NTD-targeting antibodies can facilitate the transition of the RBD to an “up” conformation, thereby enhancing the virus’s infection capacity [38]. Several NTD NAbs have been identified [26], predominantly targeting a glycan-lacking supersite [10, 27, 28, 30]. However, research on the NTD, especially following the emergence of XBB subvariants, remains inadequate. There still lacks thorough investigation into the potential of NTD-targeting antibodies to provide broad-spectrum neutralization and their underlying mechanisms.

In this study, we explored the antibody response to NTD after Omicron infection, revealing the strong immunogenicity of NTD, which elicits numerous and highly diverse antibodies targeting it. However, most of the elicited antibodies are non-neutralizing, with only a minority exhibiting neutralizing activity. These NAbs are mostly escaped by variants carrying Y144del, such as XBB.1.5. Among them, we identified five Omicron-specific antibodies that retain potent neutralizing activity against XBB.1.5 and HK.3.1. This class of antibodies exhibit a certain level of ACE2 competition and can block ACE2-mediated S1 shedding. Cryo-EM structural analysis of BD58-0730, a representative from this group, revealed that it binds to the NTD through key residues within the N1 and N2 loops. Remarkably, it interacts with the down-RBD via the light chain framework region. This class of antibodies, characterized by simultaneous interaction with NTD and RBD, exhibits ultra-potent neutralizing capabilities. This also elucidates the recent prevalent NTD mutations, such as BA.2.86 and BA.2.87.1, which carry multiple amino acid insertions or deletions in the N1/N2 loop of NTD.

Most mutations in NTD have been shown to exert minimal influence on the expression of the spike protein [39]. Also, given that SARS-CoV-2 gains entry into cells by binding to ACE2 through the RBD, mutations in the NTD are permitted within a wider range of structural tolerances. Consequently, the virus can utilize a more diverse array of mutations to counteract the immunological pressure exerted by NTD-targeting NAbs, as observed in the BA.2.86 and BA.2.87.1. Our findings provide compelling evidence suggesting that NTD represents an immunological diversion tactic employed by SARS-CoV-2. This evolutionary preference appears designed to prompt the host immune system into producing a significant quantity of either non-neutralizing antibodies or easily evadable NAbs. As a result, NTD-targeting NAbs generated from specific variants may struggle to maintain long-term efficacy. Given the flexibility of NTD, it may not represent an optimal target for vaccine design. However, within any form of vaccine design strategy based on full-length spike protein, it is crucial to evaluate the immunogenicity of NTD and the antibodies it induces, so as to circumvent the generation of deceptively high neutralizing titers.

## Materials and Methods

### Collection of plasma samples

Blood samples from individuals who had post-vaccination BA.5/BF.7 infection were collected under study protocols approved by Beijing Ditan Hospital, Capital Medical University (Ethics committee archiving No. LL-2021-024-02) and the Tianjin Municipal Health Commission, and the Ethics Committee of Tianjin First Central Hospital (Ethics committee archiving No. 2022N045KY). The information about vaccination and infection can be found in Supplementary Table 1. Blood samples were diluted 1:1 with PBS(Gibco, C10010500BT)+2% FBS (Gibco, SH30406.05) and subjected to Ficoll (Cytiva, 17-1440-03) gradient centrifugation. Plasma was collected from the upper layer and stored at -80℃ until use. Peripheral blood mononuclear cells (PBMCs) were collected at the interface and further subjected to centrifugation, red blood cell lysis (Invitrogen eBioscience, 00-4333-57), and washing steps.

### Antigen-specific cell sorting and single-cell V(D)J sequencing

PBMCs were processed with EasySep Human CD19 Positive Selection Kit II (STEMCELL, 17854) to obtain CD19^+^ B cells. B cells were stained with 3 μL FITC anti-human CD20 antibody (BioLegend, 302304, clone: 2H7), 3.5 μL Brilliant Violet 421 anti-human CD27 antibody (BioLegend, 302824, clone: O323), 2 μL PE/Cyanine7 anti-human IgM antibody (BioLegend, 314532, clone: MHM-88), 2 μL PE/Cyanine7 anti-human IgD antibody (BioLegend, 348210, clone: IA6-2), 0.013 μg biotinylated SARS-CoV-2 BA.5/BF.7 NTD protein (Sino Biological, 22BD75-77) conjugated with PE-streptavidin (BioLegend, 405204) or APC-streptavidin (BioLegend, 405207), 0.013 μg biotinylated SARS-CoV-2 WT NTD protein (Sino Biological, 40591-V49H-B) conjugated with APC-streptavidin (S3A Fig), and 0.013 μg biotinylated SARS-CoV-2 BA.5 RBD protein (Sino Biological, 22BD75-73) conjugated with PE-streptavidin. For sequencing, BA.5/BF.7 NTD was labelled with TotalSeq-C0971 streptavidin (BioLegend, 405271) and TotalSeq-C0972 streptavidin (BioLegend, 405273); WT NTD was labelled with TotalSeq-C0973 streptavidin (405275, BioLegend) and TotalSeq-C0974 streptavidin (405277, BioLegend); biotinylated Ovalbumin (Sino Biological) was labelled with TotalSeq-C0975 streptavidin (BioLegend, 405369). The cells were incubated on ice in the dark for 30 minutes. Cells were washed twice after incubation. 7-AAD (Invitrogen, 00-6993-50) was used to label dead cells. 7-AAD^−^, CD20^+^, CD27^+^, IgM^−^, IgD^−^, NTD^+^cells were sorted with a MoFlo Astrios EQ Cell Sorter. FACS data were collected by Summit 6.0 (Beckman Coulter) and analyzed using FlowJo v10.8 (BD Biosciences).

Sorted cells were processed with Chromium Next GEM Single Cell V(D)J Reagent Kits v1.1 following the manufacturer’s user guide (10x Genomics, CG000208). Cells were subjected to Gel beads-in-emulsion (GEMs) production, reverse transcription, preamplification, and V(D)J enrichment. After library preparation, libraries were sequenced with the Illumina sequencing platform.

10X Genomics V(D)J Illumina sequencing data were assembled as BCR contigs and aligned to the GRCh38 BCR reference using Cell Ranger (v6.1.1) pipeline. For quality control, only the productive contigs and B cells with one heavy chain and one light chain were kept. The germline V(D)J genes were identified and annotated using IgBlast (v1.17.1)[40]. SHM nucleotides and residues in the antibody variable domain were detected using Change-O toolkit (v1.2.0) [41].

### Expression and purification of mAbs

Antibody heavy and light chain genes were first optimized for human cell expression and synthesized by GenScript. VH and VL segments were separately inserted into plasmids (pCMV3-CH, pCMV3-CL or pCMV3-CK) through infusion (Vazyme, C112). Plasmids encoding heavy chains and light chains of antibodies were co-transfected to Expi293F™ cell (ThermoFisher, A14527) by polyethylenimine-transfection. Cells were cultured at 37℃, 8% CO_2_, 125 rpm for 6-10 days. Supernatants containing expressed mAbs were collected and further purified with Protein-A magnetic beads (Genscript, L00695).

### Preparation of SARS-CoV-2 VSV-based pseudovirus

SARS-CoV-2 spike-pseudotyped VSVs for neutralization assays were constructed as previously described using VSV pseudotyped virus (G*ΔG-VSV) [42]. Pseudoviruses carrying spikes of SARS-CoV-2 variants were constructed and used as described previously [5]. The Spike gene of SARS-CoV-2 ancestral strain (GenBank: MN908947) with specific mutations was codon-optimized for mammalian cells and inserted into the pcDNA3.1 vector. Site-directed mutagenesis PCR was performed as described previously to construct mutated sequences [43]. Briefly, variants’ spike plasmid is constructed into pcDNA3.1 vector (BA.1, A67V+HV69-70del+T95I+G142D+V143del+ Y144del+Y145del+N211del+L212I+ins214EPE+G339D+S371L+S373P+S375F+K417N+N440K +G446S+S477N+T478K+E484A+Q493R+G496S+Q498R+N501Y+Y505H+T547K+D614G+H6 55Y+N679K+P681H+N764K+D796Y+N856K+Q954H+N969K+L981F; BA.2, T19I+LPPA24-27S+G142D+V213G+G339D+S371F+S373P+S375F+T376A+D405N+R408S+K417N+N440K+ S477N+T478K+E484A+Q493R+Q498R+N501Y+Y505H+D614G+H655Y+N679K+P681H+N7 64K+D796Y+Q954H+N969K; BA.2.75, BA.2+K147E+W152R+F157L+I210V+G257S+ G339H+G446S+N460K+R493Q; BA.5, BA.2+HV69-70del+L452R+F486V+R493Q; BQ.1.1, BA.5+R346T+K444T+N460K; BQ.1.1.10, BQ.1.1+Y144del; XBB, BA.2+V83A+Y144del+ H146Q+Q183E+V213E+G339H+R346T+L368I+V445P+G446S+N460K+F486S+F490S+R 493Q; XBB.1, BA.2+V83A+Y144del+ H146Q+Q183E+V213E+G252V+G339H+R346T+ L368I+V445P+G446S+N460K+F486S+F490S+R493Q; XBB.1.5, BA.2+V83A+Y144del+ H146Q+Q183E+V213E+G339H+R346T+L368I+V445P+G446S+N460K+F486P+F490S+R493 Q; EG.5.1, XBB.1.5+F456L+Q52H; HK.3, XBB.1.5+Q52H+L455F+F456L; HK.3.1, HK.3+F157L; BA.2.86, BA.2+ins16MPLF+R21T+S50L+H69del+V70del+V127F+Y144del+ F157S+R158G+N211del+L212I+L216F+H245N+A264D+I332V+D339H+K356T+R403K+V44 5H+G446S+N450D+L452W+N460K+N481K+V483del+A484K+F486P+R493Q+E554K+A570 V+P621S+H681R+S939F+P1143L; BA.2.87.1, CVNLTTRTQLPP15-26del+A27S+HV69-70del+G75D+S98F+V126A+KNDPFLGVYYH136-146del+W152L+R190S+V213G+D215G+ G339D+S371F+S373P+S375F+T376A+D405N+R408S+K417T+N440K+K444N+V445G+L452 M+N460K+S477N+N481K+E484A+Q498R+N501Y+Y505H+D614G+P621S+V642G+H655Y+ N679R+P681H+S691P+N764K+T791I+D796H+D936G+Q954H+N969K; JN.1, BA.2+ ins16MPLF+R21T+S50L+H69del+V70del+V127F+Y144del+F157S+R158G+N211del+L212I+L 216F+H245N+A264D+I332V+D339H+K356T+R403K+V445H+G446S+N450D+L452W+L455 S+N460K+N481K+V483del+A484K+F486P+R493Q+E554K+A570V+P621S+H681R+S939F+P 1143L).

Spike-pseudotyped viruses were generated by transfection to 293T cells (ATCC, CRL-3216) with pcDNA3.1-Spike with Lipofectamine 3000 (Invitrogen). The transfected 293T cells were subsequently infected with G*ΔG-VSV (Kerafast) packaging expression cassettes for firefly luciferase instead of VSV-G in the VSV genome. Supernatants were discarded after 6-8h harvest and replaced with complete culture media. The cell was cultured for 1 day, and then the cell supernatants containing spike-pseudotyped virus were harvested, filtered (0.45-μm pore size, Millipore), aliquoted, and stored at -80 °C. Viruses of multiple variants were diluted to the same number of copies before usage.

### Pseudovirus neutralization assays

Plasma or mAbs were serially diluted and incubated with the pseudovirus in 96-well plates for 1 h at 37°C. Huh-7 cells (Japanese Collection of Research Bioresources, 0403) were seeded to the plate and cultured for 20-28 h in 5% CO_2_ at 37°C. After one-day of culture, the cells were lysed and treated with luciferase substrate (Perkinelmer, 6066769). The chemiluminescence signals were collected by PerkinElmer Ensight (PerkinElmer, HH3400).

### Enzyme-linked immunosorbent assays

SARS-CoV-2 WT NTD (Sino Biological, 40591-V49H), BA.5 NTD (Sino Biological, 22BD75-84), XBB NTD (Sino Biological, 22BD75-83) and BA.2.75 NTD (Sino Biological, 22BD75-87) were aliquoted to the 96-well plate respectively and incubated overnight at 4 ℃. The solution was removed and the plate was washed with PBST for three times. Then the plate was blocked with 3-5% BSA in PBST at 37℃ for 2h. The plate was washed for three times. 100 μL 1μg/mL antibodies were added and incubated for 30 min at room temperature, followed by five washes. Peroxidase-conjugated AffiniPure Goat Anti-Human IgG(H+L) (JACKSON, 109-035-003) was added to plates and incubated at room temperature for 15 min, followed by five washes. Tetramethylbenzidine (TMB) (Solarbio, 54827-17-7) was added and incubated for 10 min, and then the reaction was terminated with 2 M H_2_SO_4_. Absorbance was measured at 450 nm using a microplate reader (PerkinElmer, HH3400). ELISA OD450 values were normalized by dividing the highest OD450 among all dilutions, and then fitted to a two-parameter Hill function using R package mosaic (v1.8.3) to determine the midpoint titers.

### Surface plasmon resonance

Tested antibody (5 μg/mL for NTD and 1 μg/mL for spike) was captured on protein A sensor chips using a Biacore 8K (GE Healthcare). Serial dilutions of purified SARS-CoV-2 mutant spike or NTDs, ranging from 100 to 3.125 nM (for spike) or 50 to 1.5625 nM (for NTD), were injected over the sensor chips. Regeneration between concentrations is achieved with glycine 1.5. Response units were recorded at room temperature with BIAcore 8K Evaluation Software (v4.0.8.20368; GE Healthcare). The data were analyzed and fitted to a 1:1 binding model using the same software.

In the competitive binding assays, His-tagged CM5 sensor chips (Cytiva) were utilized to immobilize 5 μg/mL spike protein for 1 minute. This was followed by sequential introduction of 20 μg/mL concentrations of antibody 1 and antibody 2, each for 2 minutes. Glycine 1.5 served as the regeneration agent. The SPR competition score for *a* as first antibody and *b* as second antibody is defined as:

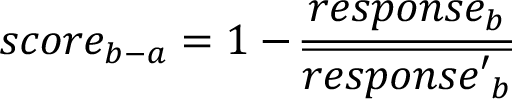

In the formula, *response_b_* represents the response units when *b* serves as the second antibody and *a* as the first antibody, whereas 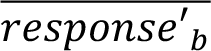 denotes the mean response units when *b* acts as the first antibody.

### S1 shedding

The cell supernatants containing pseudovirus particles were incubated with soluble huACE2-Fc at different concentrations at 37℃ for 1 h. Afterward, virus particles were pelleted at 18,000 xg for 1 h at 4℃. The pelleted virus particles were washed once with cold PBS before the samples were resuspended in 1 X Sample buffer (Solarbio, P1016). Then, samples were subjected to SDS-PAGE and transferred to NC membrane (Cytiva, A29696255) for western blotting analysis. The membranes were blocked and incubated with primary antibodies at 37℃ for 1 h to detect the S1 and S2 subunits using rabbit anti-SARS-Spike S1 (Sino Biological, 40591-T62) and rabbit anti-SARS-Spike S2 (Sino Biological, 40590-T62) respectively. After washing, goat anti-Rabbit IgG/HRP (Solarbio, SE134) was incubated at room temperature for 30 min. Finally, chemiluminescent substrate (Thermo, 34577) was added before the membranes were analysed with ChemiDoc System (Bio-Rad).

### MACS-based antibody-escape mutation profiling

Duplicated DMS libraries were constructed based on BA.5 NTD (position S13-K304 by Wuhan-Hu-1 reference numbering) as previously described [5]. Briefly, site-directed mutagenesis PCR was performed using rationally designed NNS primers to create all 5740 possible amino acid mutations. Then each NTD variant was PCR appended to a unique 26-neuclotide (N26) barcode, and the link of N26 barcode and NTD variant was obtained by PacBio sequencing. Two BA.5 NTD DMS libraries were further assembled into pETcon 2649 vector (Addgene) and electroporated into electrocompetent DH10B cells for plasmid amplification, then transformed into the *Saccharomyces cerevisiae* EBY100 strain. The obtained yeast population was screened on SD-CAA plates and further enlarged in SD-CAA liquid media.

The yeast libraries were first inoculated into SG-CAA media to induce the expression of NTD variants and the properly expressed NTD variants were selected by fluorescence-activated cell sorting (FACS) and further prepared for antibody mutation escape profiling. In order to capture NTD variants that escape BD58-0730 binding, two rounds of negative selection by Protein A beads (Thermo Fisher) and one round of positive selection by anti-c-myc antibody conjugated beads (Thermo Fisher) were performed as previously reported [5]. The obtained yeast population was overnight amplified and collected for plasmid extraction using Zymoprep™ Yeast Plasmid Miniprep II (ZYMO RESEARCH). The N26 barcode was PCR amplified using the above extracted plasmid as the template and final PCR products were purified by Ampure XP beads (Beckman Coulter), quantified, and sequenced on Illumina Nextseq 550 platform.

### DMS data analysis

The raw DMS sequencing data were processed as previously described [5, 6]. Briefly, the barcode sequences detected in both the antibody-screened and reference library were aligned to the barcode-variant dictionary generated from PacBio sequencing data of the BA.5 DMS library using dms_variants (v0.8.9). To avoid large sampling errors, only barcodes detected more than 5 times in the reference library were included in the analysis. The escape scores for a variant X detected in both the screened and reference libraries were defined as F × (nX,ab/Nab)/(nX,ref/Nref), where F is a scale factor normalizing the scores to the 0–1 range, and n and N represent the number of detected barcodes for variant X and the total barcodes in the antibody-screened (ab) or reference (ref) samples, respectively. To assign an escape score to each single substitution on the NTD, an epistasis model was fitted using dms_variants (v0.8.9) as previously described [44, 45]. For antibodies with multiple replicates of DMS, the final escape score for each mutation was the average over all replicates.

### Protein expression and purification for structure analyses

HEK293F cells were maintained at 37 ℃ and 5% CO2 in SMM 293-TI medium (Sino Biological) supplemented with penicillin-Streptomycin. The full length BD58-0730 and XBB.1.5 S6P were obtained by transient expression in the HEK293F cells using polyethylenimine (PEI, Polyscience). Culture supernatants were harvested at 96 h post transfection, exchanged into the binding buffer (25 mM Tris-HCl, pH 7.4, 150 mM NaCl), and proteins were purified from the conditioned media using the Ni-NTA and size exclusion chromatography. The final buffer used for the gel filtration step contains 20 mM HEPES, pH 7.4, and 150 mM NaCl.

### Cryo-EM data collection, processing, and structure building

To prepare the sample for Cryo-EM, 1 mg BD58-0730 and 1 mg S6P were mixed on ice for an hour, and separated antibody-antigen complex using size exclusion chromatography. The complex (1 mg/mL) was then applied onto the glow-discharged holy-carbon gold grids (Quantifoil, R1.2/1.3) using an FEI Vitrobot IV. The grids were blotted with a filter paper at 4 ℃ and 100% humidity, and flash-cooled in liquid ethane and screened using a 200 KV Talos Aectica. Data collection was carried out using a Titan Krios electron Microscope operated at 300 KV. Movies were recorded on a K3 summit direct electron detector using the EPU software. A total of 3046 movies were recorded for image processing. A total dose of 50 e-/ Å2 was accumulated on each movie comprising 32 frames with a pixel size of 0.54 Å and a defocus range of -3.00 μm.

CTF estimation, particle-picking, extraction, 2D classification, 3D classification, homogeneous refinement and local refinement were carried out using cryoSPARC. The workflow of data processing was illustrated in S6 Fig. The S trimer (PDB: 8IOT) and the Fv region of BD58-0730 predicted by Swiss model were docked into the cryo-EM density map using UCSF Chimera. The structure models were then manually built in coot and refined using the real-space refinement in PHENIX. Figures were prepared using UCSF Chimera.

To elucidate the mechanism by which BD58-0730 competitively inhibits the binding of S protein to ACE2, the RBD-ACE2 (PDB: 7C8D) complex was aligned with the “up” RBD of the S-BD58-0730 complex and the minimal distance between ACE2 and BD58-0730 light chain was measured. The light chain of BD58-0730 bound to NTD was predicted by the Swiss model.

## Acknowledgments

We thank all volunteers for providing blood samples.

## Supplemental information

**S1 Table. Summarized information of SARS-CoV-2 convalescents, related to Fig 1 and Fig 4.**

**S2 Table. Neutralizing activities, VDJ genes and CDR3 sequences of 361 NTD antibodies.**

**S1 Fig. Isolation and characterization of NTD antibodies.** (A) Schematic of the SARS-CoV-2-related immune histories of the cohorts involved in this study. (B) FACS strategy for sorting memory B cells encoding NTD-targeting antibodies from PBMCs. Percentage of cells in each gate are labeled. Representative results of multiple batches are shown. (C) heatmap of neutralizing activities against pseudovirus and ELISA OD450against WT, BA.2.75, BA.5 and XBB NTD of 361 NTD-targeting mAbs.

**S2 Fig. Binding affinity of 5 ultra-potent NAbs defined by SPR.** (A) SPR sensorgrams of 5 ultra-potent NAbs against BA.5 NT(D) XBB NTD and XBB.1.5 spike. (B) ka, k(D) and KD results of (A).

**S3 Fig. Sequence and ACE2 competition analysis of 5 ultra-potent NAbs.** (A) CDR3 sequence similarity of 5 ultra-potent NAbs and 10 randomly selected NTD antibodies from the same batch. (B) Pseudovirus-neutralizing IC50 of antibodies in Fig. 2b against SARS-CoV-2 D614G, BA.1, BA.2, BA.2.75, BA.5, BQ.1.1, BQ.1.1+Y144del, XBB, XBB.1, XBB.1.5, EG.5.1 and HK.3.1, and their usage of heavy chain V(VH), heavy chain J (VJ), light chain V (VL) and light chain J (VJ) genes. (C) SPR competition scores of Fab and F(ab’)2 form of 5 ultra-potent NTD-targeting NAbs, BD57-3253 and BD55-1205 as first antibody against ACE2.

**S4 Fig. Blockage of ACE2-induced S1 shedding by 5 ultra-potent NAbs.** (A) S1 shedding assay of 5 ultra-potent NAbs against BA.5 and XBB.1.5. BA.5 and XBB.1.5 pseudovirus particles were pre-incubated with the 5 antibodies for 1 hour before a subsequent 1-hour incubation with 5μg/mL hACE2-Fc. Western blot analysis was employed to assess the retention of S1 and S2 subunits.

**S5 Fig. Workflow for BD58-0730 Cryo-EM reconstructions.** (A) A representative raw image collected using a Titan Krios300 kV microscope with a K3 detector. Representative 2D classes. Gold standard Fourier shell correlation (FSC) curve with estimated resolution.Viewing Direction Distribution Iteration. (B) Flow chart of image processing.

**S6 Fig. Pseudovirus-neutralizing assay of 5 ultra -potent NTD-targeting NAbs against XBB.1.5 and XBB.1.5+N481K.** (A) Pseudovirus-neutralization curve of 5 ultral-potent NAbs against XBB.1.5+N481K. Data for each mAb were obtained from a representative neutralization experiment, which contains four replicates. Data are represented as mean ± SD. (B) Pseudovirus-neutralizing IC50 of 5 ultra-potent NTD-targeting NAbs against XBB.1.5 and XBB.1.5+N481K.

